# Phasor EO-FLIM: Lifetime imaging with picosecond noise and 500 Hz frame rate

**DOI:** 10.64898/2026.05.29.728614

**Authors:** Linghao Hu, Pingchuan Ma, Vaibhav Menon, Athira Sethu Madhavan, Kenta Asahina, Lena M. Müller, Adam J. Bowman

## Abstract

Fluorescence lifetime microscopy is limited by photon throughput, constraining the speed and dynamic range of biological measurements. We demonstrate a compact optical module and fast phasor acquisition methods to image multi-exponential lifetimes at 500 Hz *in vivo* and with over **10^10^** photons per second in static tissues. Action potentials are captured in phasor plots, and lifetime contrast is revealed from autofluorescence and wheat germ agglutinin stain with 20 picosecond pixel noise.

---

Fluorescence lifetime imaging (FLIM) is a powerful method for achieving quantitative measurements from genetically encoded fluorescent sensors, chemical probes, and endogenous autofluorescence. Analysis of multi-component decays is central to lifetime imaging and often uses polar phasor plots to represent each pixel’s decay without explicit fitting [1]. Despite the utility of existing multi-exponential and phasor FLIM approaches, they remain limited by photon acquisition rates or the performance of gated and modulated camera sensors [2–4]. Frame acquisition rates are generally less than 10 Hz, and lifetime sensitivity is fundamentally limited by the acquired number of photons per pixel [5–8].

We enabled fast wide-field phasor FLIM by engineering compact radio frequency optical gating modules and developing multi-phase acquisition and analysis methods. Electro-optic FLIM (EO-FLIM) is an emerging technique that enables wide-field FLIM using polarization modulation to gate fluorescence into two images on a scientific camera [9]. Initial demonstrations focused on single-exponential estimation using a fixed phase offset between laser pulses and the optical gate and have faced technical limitations due to thermal dissipation and bulky high-voltage drivers [10–13]. Here, we designed radio frequency EO-FLIM modules based on improved electro-optic crystal mounts and planar resonant circuits (Fig. 1a and Extended Data Fig. 1a). The modules are compact (2.5 × 2.5 × 5 inches) (Extended Data Fig. 1c,d), remove limitations of crystal thermal expansion, and are easily integrated into any microscope. We developed multi-exponential phasor analysis and triggering methods to execute a table of programmed phase offsets synchronously with the camera shutter, enabling fast phasor FLIM acquisition at frame rates of up to 500 Hz *in vivo*, corresponding to 2500 Hz sampling of individual gated images.

**Fig. 1.**
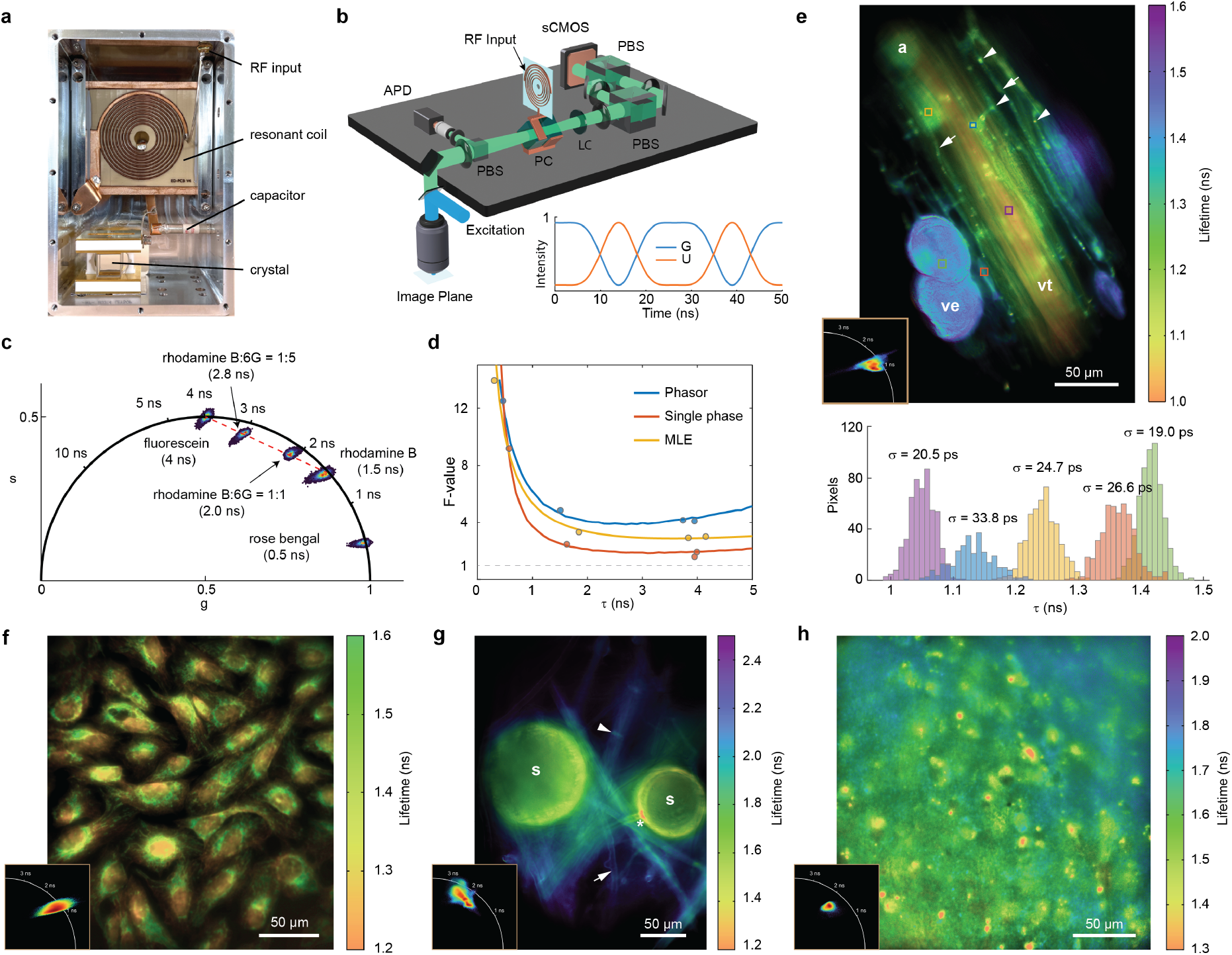
Wide-field phasor EO-FLIM. (**a**) Pockels cell (PC) and planar resonant circuit module achieves 40 MHz gating across a 300 *µm* × 300 *µm* FOV (Extended Data Fig. 1 and Supplementary Fig. 1d,e). (**b**) System schematic and instrument response function. Fluorescence is gated by the PC at 40 MHz. Gated (*G*) and ungated (*U*) channels are separated using polarizing beam splitters (PBS) and separately focused onto an sCMOS camera. An avalanche photodiode (APD) samples a small fraction of the fluorescence. A liquid crystal waveplate (LC) cancels residual birefringence. (**c**) Phasor plot of fluorescent dyes in solution. (**d**) Simulated and experimental F-values comparing single-phase, phasor, and maximum likelihood estimation. Data points for fluorescein (*τ* = 4 ns), rhodamine B (*τ* = 1.5 ns), rhodamine 6G (*τ* = 3.95 ns), and rose bengal (*τ* = 0.5 ns) solutions are shown. **(e-f)** Phase lifetime images and inset phasor plots. (**e**) *Medicago truncatula* root colonized by symbiotic *Rhizophagus irregularis* fungi and stained with Alexa Fluor 488 WGA (1.1 × 10^11^ photons in 14.5 s). Lifetime histograms and standard deviations are shown for selected ROIs. Plant vascular tissue (vt) and fungal arbuscules (a), hyphae (arrows), septa (arrowheads), and vesicles (ve) within the root cortex are distinguished. (**f**) Fixed U2OS cells stained with Alexa Fluor 532 (mitochondria) and Alexa Fluor 555 (microtubules) (1.8 × 10^11^ photons in 29 s). (**g**) Autofluorescence from *Rhizophagus irregularis* fungal spores (2.2 × 10^11^ photons in 14.5 seconds). Spores (s), hyphae (arrow), septa (arrowhead), and germ pore (asterisk) are visible. (**h**) Autofluorescence from an acute mouse brain slice (7.7 × 10^10^ photons in 14.5 s).

We integrated our EO-FLIM module into a custom upright epifluorescence microscope (Fig. 1b, Supplementary Fig.1a, and Methods). The system allows both single-phase and multi-phase EO-FLIM acquisition. Single-phase acquisition samples the convolution of the fluorescence decay with the gating function at a single point, relating average lifetime to intensity ratio through a lookup table as previously described (Supplementary Fig. 2a,b) [9, 10]. For phasor acquisition, we sample the convolution of the resonant gate with the fluorescence decay at multiple phase points and construct the phasor coordinates for each image pixel by discrete Fourier transform and deconvolution with the instrument response function (IRF) (Supplementary Fig. 2c,e and Methods). Phases are advanced synchronously with the camera shutter in a continuous loop (Supplementary Fig. 3). A time-correlated single photon counting (TC-SPC) detector samples a fraction of the collected photons to provide spatially-averaged ground truth data.

We first validated system performance <u>us</u>ing fluorescent dye samples and quantified the lifetime sensitivity or F-value 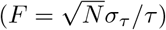 for single-phase, phasor, and maximum likelihood estimation (MLE) (Fig. 1c,d). Phasor estimation achieves an F-value of 3.86 compared to 1.84 for single-phase estimation. We characterized the performance of phasor estimation with different sub-sampling strategies. The phase lifetime F-value is independent of the number of sampled phases, and we can achieve reliable imaging with as few as five sampled points (Supplementary Figs. 4 and 5). In contrast to single-phase estimation, the phase lifetime is robust to wavelength-dependent and spatially-dependent gating effects in the crystal, allowing precise lifetime imaging across large FOVs (Supplementary Fig. 6d,e and Supplementary Derivation). Spatial dependence of the modulation lifetime can be corrected through low-pass image filtering and comparison to TC-SPC data as needed (Supplementary Fig. 7). By sampling five phase points, we readily captured fast phasor dynamics at 200 Hz while mixing rhodamine B and 6G dye solutions and 500 Hz when quenching fluorescein dye with potassium iodide (Extended Data Fig. 2b,c and Supplementary Videos 1 and 2).

In static tissues, phasor EO-FLIM achieves high dynamic range and picosecond noise. We imaged Alexa Fluor 488 wheat germ agglutinin (WGA) conjugate, a common stain for cell membrane glycans and chitin, in cleared samples of *Medicago truncatula* roots colonized by three symbiotic arbuscular mycorrhizal fungi (AMF) species (see Methods). We found that the stain displays structurally-specific lifetime contrast within a 500 picosecond dynamic range, allowing discrimination of AMF fungal structures such as arbuscules, vesicles, and hyphae as well as root tissue layers (Fig. 1e and Extended Data Fig. 3). This contrast likely reflects local changes in fungal cell wall structure and composition [14, 15]. EO-FLIM phasor estimation is in good agreement with results from TC-SPC, single-phase, and fitting analysis (Supplementary Fig. 8). Fixed U2OS cells (GATTAquant GmbH) were used to demonstrate phasor FLIM imaging of mitochondria and microtubules, resolving Alexa Fluor 532 and Alexa Fluor 555 labels within the same spectral channel (Fig. 1f). We also acquired and segmented images based on their autofluorescence phasor profiles, as demonstrated with *Rhizophagus irregularis* spores and acute brain slices where we observed rich lifetime contrast (Fig.1g,h and Extended Data Fig. 4). We detected 2.2 × 10^11^ photons when capturing 145 phase points in 14.5 seconds (Fig. 1g), which would require over two days to acquire at 1 MHz counting rate in a single photon counting system. Local precisions and pixel noise levels *∼*20 picoseconds were achieved (Fig. 1e and Extended Data Fig. 5b,c).

We performed dynamic phasor FLIM imaging *in vivo* with large fields of view and acquisition rates up to 500 Hz. We first demonstrated 40 Hz phasor imaging of a crawling *Drosophila melanogaster* larva labeled pan-neuronally with a green fluorescent protein. Autofluorescence from tissue, gut, and labeled neurons was clearly visualized (Fig. 2a,b and Supplementary Video 3). Phasor FLIM imaging at 200 Hz captured sub-threshold voltage transitions in a single neuron expressing the genetically encoded voltage indicator pAce [16] in adult *Drosophila* (Fig. 2c-e and Supplementary Video 4). Voltage dynamics over time are represented by the trajectory of the mean phasor coordinate (Fig. 2f and Supplementary Video 4b), and a lifetime difference of approximately 100 picoseconds between depolarized and hyperpolarized states is resolved in the phasor plot (Fig. 2g). At smaller fields of view, we captured action potentials with 500 Hz phasor acquisition rate (Fig. 2h and Supplementary Video 5). The acquired field of view for fast phasor imaging is limited only by the camera’s rolling shutter speed.

**Fig. 2.**
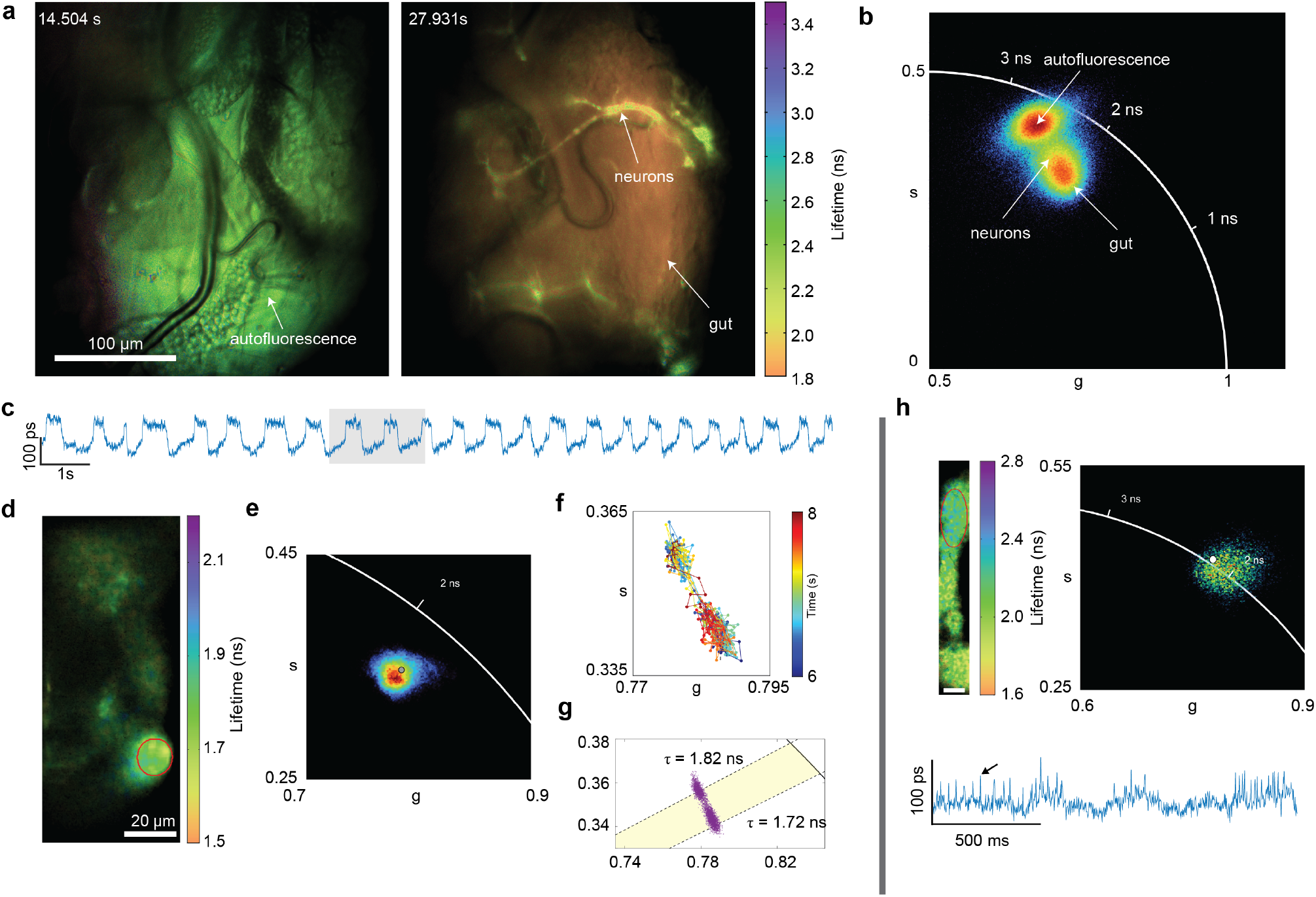
Fast phasor EO-FLIM *in vivo*. (**a**) 40 Hz FLIM of a crawling *Drosophila* larva (Supplementary Video 3). (**b**) Phasor plots corresponding to the two displayed frames overlaid. **(c-g)** 200 Hz phase lifetime recording of sub-threshold voltage in a *Drosophila* neuron (Supplementary Video 4). **(c)** Phase lifetime recording from ROI shown in image (**d**). (**e**) Corresponding phasor point cloud acquired at 200 Hz. The gray dot represents the ROI average coordinate. (**f**) Time trajectory of mean phasor coordinates for the shaded times in (**c**). (**g**) Scatter plot of mean coordinate showing sub-threshold states with 100 ps lifetime difference. (**h**) 500 Hz phasor imaging of a spiking neuron in *Drosophila* (left, scale bar 8 *µm*), a single phasor plot acquired at 500 Hz (right), and the phase lifetime recording showing action potentials (bottom − arrow indicates representative action potential). All images report phase lifetime.

Phasor EO-FLIM and compact planar resonant Pockels cells are broadly compatible with any wide-field microscope, allowing efficient lifetime imaging with low noise, picosecond resolution, and acquisition rates up to 500 Hz. We anticipate application to quantitative readout of fluorescent biosensors and profiling of tissue autofluorescence as well as future extensions to faster sampling and spectral-lifetime multiplexing.

## Methods

### Optical design

Experiments were performed on a custom upright epifluorescence microscope optimized for *in vivo* imaging (Fig. 1b). The system was designed based on Zemax OpticStudio (Ansys) simulations to achieve *∼*300 *µ*m field of view when using a 20X objective (Supplementary Fig. 1a). The sample is excited with a 40 MHz supercontinuum laser (YSL, SC-Pro) supplying *∼*100 picosecond excitation pulses. The excitation band (400 nm - 700 nm) is controlled using motorized linear bandpass filters (YSL, VLFM3580). The laser passes through a variable beam expander before entering a fluorescence filter cube assembly (Thorlabs WFA2002, TLV-U-MF2-BFP). Either 20X, 0.95 NA, 2 mm working distance (Olympus XLUMPlanFI) or 16X, 0.8 NA, 3 mm working distance (Nikon MRP07220) water immersion objectives are used. The Pockels cell is located at the first image plane, allowing the position of the crystal to be adjusted to optimize gating performance (Supplementary Fig. 1a). This image is then relayed to a Photometrics Kinetix scientific CMOS camera (10.2 megapixels, 29.4 mm diagonal) using a 4f system (Thorlabs MPTTL200 and TTL200) to capture the gated and ungated images. For all presented data, we operate the camera in dynamic range mode with a maximum full frame rate of 83 Hz and 1.6 electrons/pixel read noise.

The gated (*G*) and ungated (*U*) images are split and recombined on the camera sensor using a pair of 40 mm broadband polarizing beamsplitter cubes (Bernhard Halle Nachfl. GmbH) with intermediate lenses (Thorlabs TTL200) to control image focus (Supplementary Fig. 1b). A fraction of the fluorescence is directed to a diffuser and focused onto an avalanche photodiode (MPD $PD-050-CTD) with a TC-SPC detector (Swabian Time Tagger Ultra), providing spatially average ground truth data. A liquid-crystal variable waveplate (Meadowlark LVR-200-VIS) positioned after the Pockels cell compensates for static birefringence.

### Pockel cell and radio frequency electronics

Imaging Pockels cells were designed using pairs of lithium tantalate crystals with transverse electric field and broadband visible anti-reflection coating (Leysop Inc.). To avoid thermal expansion limitations on modulation depth and field of view – a significant limitation in earlier prototype devices [10, 11, 13] – we developed two mounting schemes that gave similar optical gating performance and stability across drive powers (Extended Data Fig. 1c,d and Supplementary Fig. 1d-g). The first configuration (Extended Data Fig. 1c) splits the top crystal electrodes using a flexible copper foil bridge, and the second (Extended Data Fig. 1d) uses an achromatic half wave plate between crystals so that the crystal axes are perfectly aligned for each pair, matching thermal expansion coefficients. Both configurations are compatible with pairs of 10×10×10 mm and 15×15×10 mm crystals. We expect future use of higher quality crystal material can further improve spatial uniformity.

Planar resonant transformers were designed on low loss ceramic circuit boards (aluminum oxide or aluminum nitride) with a primary coupling loop impedance matched to 50 Ω drive electronics and tuned to achieve a 20 MHz resonant frequency with the PC. Both the PC bottom electrode and transformer board edges are heat sunk to a water-cooled aluminum housing. Fine frequency tuning is provided by either a trim capacitor or an adjustable copper plunger located behind the planar coil (Fig. 1a and Extended Data Fig. 1b).

The laser provides a common 40 MHz clock to a direct digital synthesizer (DDS - Novatech 409B-AC) and the TC-SPC electronics (Supplementary Fig. 2). The optical gating waveform has twice the frequency of the radio frequency drive. The DDS generates a phase-locked 20 MHz drive signal with a programmable phase offset. The drive signal is amplified using a pre-amplifier (Mini-circuits, ZHL-42) and a class-AB amplifier (Empower, 1058-071), while transmitted and reflected power are monitored using a SWR meter (Nissei RS-70). Drive powers ranging from 25 to 60 Watts are used.

For an applied resonant drive voltage amplitude *V* at frequency *ω*, the Pockels cell generates a time-dependent birefringent phase shift, resulting in an intensity modulation in the gated and ungated channels through crossed polarizers:

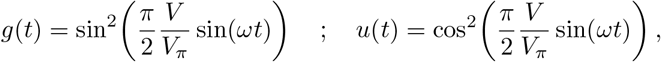

where *V*_*π*_ is the half-wave voltage, which is approximately 1.2 kV for 10 mm and 1.8 kV for 15 mm crystals at 550 nm.

### IRF measurement

The IRF or gating function was measured using dye solutions of quenched fluorescein or methylene blue in a fluidic channel slide (Ibidi µ-Slide I Luer, 80166). An affine registration was applied to register the *G* and *U* images using an image of a stained *Convallaria* rhizome slide (Boston Electronics) or a 10 micron grid (Electron Microscopy Sciences, slide S29).

### Lifetime estimation

The normalized gated and ungated intensities are determined by the convolution of IRF with the fluorescence decay 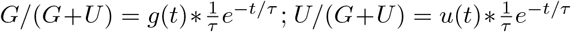. For single-phase estimation, the optimal phase is chosen from the IRF (Supplementary Fig. 2a), so that the measured *G/U* ratio is maximally sensitive to the estimated lifetime. A lookup table is then generated to map fluorescence lifetime to the corresponding G/U ratio for the selected phase offset (Supplementary Fig. 2b). If multiple phase points of the intensity trace are acquired (Supplementary Fig. 2c), the fluorescence lifetime can be estimated using either phasor analysis, least squares trace fitting, or maximum likelihood estimation (MLE) (Supplementary Fig. 2d,e).

To perform phasor FLIM acquisition, the *G* and *U* intensities are measured at multiple phase points spanning at least one full gating cycle, including both the starting (0°) and ending (180°) phase points to compensate for rolling shutter phase offsets. A minimum of five phase points within the phase sweep (0°, 45°, 90°, 135°, and 180°) is required to maintain estimation precision. The phase lifetime F-value is independent of the number of sampled points, accounting for photon division between frames (Supplementary Fig. 5c). Phasor analysis can be performed either on the normalized gated intensity *G/*(*G* + *U*) or gated intensities *G* alone by calculating the first-harmonic Fourier component discretely over N sampled phases as

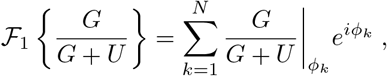

where *ϕ*_*k*_ is the *k*-th sampled phase offset. The IRF is then deconvolved by

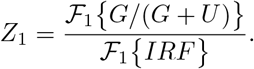

The phasor coordinates g (x-coordinate) and s (y-coordinate) for each pixel correspond to the real and imaginary components of *Z*_1_ respectively. The phase and modulation lifetime estimates can then be calculated from g and s (see Supplementary Derivation).

### Phasor plots

#### Acquisition

Our standard phasor EO-FLIM acquisitions captured 145 frames at phase offsets from 0° to 360° in 2.5° increments, covering two gating cycles. Frame exposure times of 100 or 200 ms were used, depending on the sample, resulting in total acquisition times of 14.5 or 29 s. Fast phasor EO-FLIM was performed by capturing five phase points: 0°, 45°, 90°, 135°, and 180° (Fig. 2 and Supplementary Videos 3 - 5). Depending on the sample and the field of view, frame exposure times were as short as 400 *µ*s, enabling phasor imaging up to 500 Hz. For imaging of fluorescein solution (Fig. 1c), *Medicago truncatula* roots (Fig. 1e), *Rhizophagus irregularis* fungal spores (Fig. 1g), acute brain slices (Fig. 1h), and *Drosophila* (Fig. 2) samples were excited at 450 - 490 nm with emission collected at 500 - 550 nm. Rhodamine and rose bengal dye solutions (Fig. 1c) were excited at 520 - 550 nm with emission collected at 570 - 610 nm. Fixed U2OS cells (Fig. 1f) were excited at 490 - 520 nm with emission collected above 532 nm. All lifetime images were acquired using the 20× objective, except for the *Drosophila* larva shown in Fig. 2a,b and Supplementary Video 3, which was acquired using the 16 × objective. The laser power was between 6 to 18 mW depending on the sample.

#### Display

For standard phasor FLIM imaging, all phasor plots were generated from the normalized gated intensity trace *G/*(*G* + *U*) with pixelwise IRF deconvolution, except for the phasor plot in Fig. 1e,g, which was generated from the raw gated intensity trace (*G*) and global IRF. Median filtering or spatial binning was applied to the modulation for selected datasets (Fig. 1e-g and Extended Data Fig. 3). Finally, phase and modulation calibration were performed based on the global TC-SPC point (Fig. 1e-h and Extended Data Fig. 3). For fast phasor FLIM imaging, only modulation calibration was applied to selected datasets using TC-SPC coordinates calculated from either a 5 frame temporal window (Fig. 2b and Supplementary Video 3) or the entire recording (Fig. 2e and Supplementary Video 4). At short frame exposure times (<10 ms), the rolling shutter of the sCMOS camera can introduce a phase shift along the vertical axis, resulting in a systematic lifetime gradient. This effect was corrected by applying a planar correction to the lifetime image, referenced to the middle row. In Fig. 2e,h, the *g* and *s* images were denoised using a Gaussian filter of *σ* = 5 and *σ* = 3 pixels, respectively. 2×2 and 4×4 binning were used for phasor plot of Fig. 2b and Supplementary Video 3 to improve visualization. Both color and transparency of the phasor points were defined by local point density, except for Supplementary Video 3 (intensity). Lifetime images were displayed by combining the phase lifetime image with a transparency mask generated from intensity. A Gaussian filter can be applied to flatten the intensity image when generating the mask.

#### Calibration

Both the phase and modulation lifetimes measured by EO-FLIM can be calibrated using the spatially averaged TC-SPC data by applying a global correction. Every captured image file is saved along with a simultaneous TC-SPC histogram (10 ps bin width) (Supplementary Fig. 3). Phase angle deviation primarily results from discrete sampling (Supplementary Fig. 5a-e) or IRF mismatch, while modulation deviation is due to spatially non-uniform gating within the crystal (Supplementary Fig. 1d-g) and wavelength-dependent effects (Supplementary Fig. 6d,e, and Supplementary Derivation). Typical global phase angle deviations are small if present (< 1° or < 100 picoseconds), so local modulation corrections are more dominant (Supplementary Fig. 7). Because modulation deviations occur at low spatial frequencies in the crystal, they can be separated from image structures by filtering.

To perform phase shift calibration, the intensity-weighted average (IWA) of the g and s coordinates is calculated. The phase offset between the IWA and the TC-SPC reference Δ*θ* is applied to all pixels as a rotation in the phasor plot, while preserving each pixel’s radial distance from the origin (modulation depth) (Supplementary Fig. 7a). For modulation calibration, a Gaussian filter is applied to the modulation image to extract its low-frequency component, which represents the spatially varying background modulation (Supplementary Fig. 7b). The standard deviation of the Gaussian filter is adjusted according to image size, with *σ* typically from 50 to 100 pixels. The difference between the background modulation and the TC-SPC modulation (Δ*m*) is used to generate a spatial modulation calibration that is applied to all pixels (Supplementary Fig. 7b). Finally, the g and s coordinates are recalculated, preserving the original phase angle for each pixel.

#### Δ*τ* calculation

To quantify the lifetime dynamics of *Drosophila* neurons over time within a fixed ROI, intensity frames were generated by summing raw images acquired at every five phase points (0°, 45°,90°, 135°, 180°). Sample motion was corrected using piecewise affine registrations calculated every 100 intensity frames. After correction of the rolling shutter effect and filtering pixel noise, lifetime images were computed from the phase angle of the resulting *g* and *s* coordinates. The mean *g, s*, intensity, and lifetime were calculated within the ROI for each time point. Δ*τ* was obtained by subtracting a 500 point moving average from the lifetime trace. A Gaussian filter with *σ* = 10 pixels was applied to the Δ*τ* image in Supplementary Video 4a for visualization.

### F-value simulations

Monte Carlo simulations were performed to evaluate the photon efficiency of EO-FLIM. The experimentally measured IRF was convolved with single-exponential decays to generate phase traces. The simulation was repeated 1000 times to calculate F-values using 10^11^ photons distributed across 145 evenly spaced phase points with Poisson statistics. The fluorescence lifetime was estimated using single-phase, phasor analysis, and maximum likelihood estimation (MLE) methods, and F-values were calculated for each. MLE was performed by minimizing the negative Poisson log-likelihood of the observed G and U photon counts for a single-exponential decay using the integrated expected counts computed at each sampled gate phase. For comparison to experimental data (Fig. 1d), the experimental F-value was calculated from the phase lifetime using the central 200 × 200 image pixels.

### Sample preparation

#### Dye solutions

Fluorescein (700 *µM*), rhodamine B (100 *µM*), rhodamine 6G (100 *µM*), and methylene blue (100 *µ*M) were prepared in water, while rose bengal (1 *mM*) was prepared in methanol. The fluorescein dye was quenched by mixing 170 *µL* saturated fluorescein with 1 *mL* saturated potassium iodide.

#### Acute brain slice

Coronal sections from a naive adult mouse brain (300 *µ*m thick) were prepared with a vibratome (Leica Instruments, VT1200S) in cold sucrose-based cutting solution (concentrations in mM: 87 NaCl, 2.5 KCl, 25 NaHCO_3_, 1.25 NaH_2_PO_4_, 1 MgCl_2_, 0.5 CaCl_2_, 25 glucose, and 75 sucrose). After sectioning, slices were transferred to artificial cerebrospinal fluid (ACSF) solution (concentrations in mM: 127 NaCl, 2.5 KCl, 25 NaHCO_3_, 1.25 NaH_2_PO_4_, 1 MgCl_2_, 2 CaCl_2_, and 25 glucose). Following incubation at 34 °C for 10 min, the slices were kept at room temperature. For imaging, the slices were transferred to a microscope chamber, with ACSF perfused at a flow rate of 2 - 4 mL/min. All solutions were continuously bubbled with 5% CO_2_ and 95% O_2_.

#### *Medicago truncatula* roots/AMF

*Medicago truncatula* roots were inoculated with and stained for the AMF *Rhizophagus irregularis* (DAOM197198), *Gigaspora rosea* (INVAM WV187-12), or *Paraglomus occultum* (INVAM VA102A-24) as previously described [17]. In brief, 5-day-old *M. truncatula* seedlings were transplanted into SC10 cone-tainers (Steuwe and Sons) filled with a 1:1 mixture of sterile play sand and fine vermiculite. For *R. irregularis* inoculation, 250 spores (Premiertech, Canada) were placed 5 cm below the substrate surface. For inoculation with other AMF species, 5 g of *G. rosea* and *P. occultum* inoculums (consisting of spores, hyphae, roots and substrate from a 2 months-old bahia grass-AMF preculture) were mixed thoroughly into 100 g of sterile 1:1 play sand:vermiculite substrate. Plants were co-cultivated with AMF for 5 weeks in a growth chamber set to 16 h light (24°C)/8 h dark (22°C) and 40% relative humidity, during which they received biweekly treatment with 15 ml of 0.5× Hoagland’s nutrient solution supplemented with 20 µM phosphate to encourage symbiosis development. At the end of the experiment, roots were extracted from the substrate, washed in ddH2O, and cleared in 20% KOH for 3 days at 65°C. Roots were then rinsed three times in ddH2O, followed by overnight incubation in 0.2 mg ml^−1^ WGA-Alexafluor488 (Thermo Fisher Scientific) in 1× PBS to stain fungal structures.

For AMF spore imaging, *R. irregularis* spores were resuspended in ddH2O and allowed to germinate at room temperature for 7 days. Germinated spores were directly imaged in ddH2O solution without additional staining.

#### Drosophila

*Drosophila* larvae and adults expressing the genetically encoded voltage indicator pAce were imaged. For larvae, this indicator provided structural pan-neuronal labeling using the elav-GAL4 line. To image crawling, second instar larvae were placed in a 200 *µ*m-wide channel filled with 2% agarose using a 0.3 mm thick plastic film spacer between No. 1.5 coverslips. Neuron activity in adult flies carrying × 20 UAS-pAce [16] and either MB085C-GAL4 (MBON-*γ*1pedc*> α/β*, Fig. 2c - g) or MB504B-GAL4 (PPL1 dopaminergic neurons, Fig. 2h) [18] was imaged. Five-week-old flies were mounted on a custom holder, and an imaging window was created in the cuticle using a tungsten needle (Fine Science Tools, 10130-05) or hypodermic needle (BD Precision Glide, 27 Gauge). After surgery, *∼*1 *µ*L of UV-curable optical adhesive (NOA68, Norland) was applied, and a small piece of #0 coverglass was placed over the opening to seal the window.

## Supplementary information

This article is accompanied by 5 Extended Data Figures, 8 Supplementary Figures, and 5 Supplementary Video Files.

## Acknowledgements

We thank Cheng Huang for providing the pAce fly lines and Rebeca Osca-Verdegal for assistance with brain slice preparation. This work was supported in part by funding from the Biohub Imaging Residency Program, the Dan and Martina Lewis Biophotonics Fellowship (L.H.), the Pioneer Fund Postdoctoral Scholar Award (P.M.), USDA-NIFA award 2022-67013-42820 (L.M.M.), NIH T32 NS136094 (V.M.), and NIH R35 GM119844 (K.A.).

## Author Contributions

L.H. and A.J.B. designed and built the microscope, developed phasor methods, performed imaging experiments, and wrote the manuscript. L.H., P.M., and A.J.B. performed data analysis. P.M. performed brain slice imaging. A.S.M. and L.M.M. provided *Medicago truncatula* root and AMF samples. V.M. and K.A. provided surgically prepared *Drosophila*. All authors contributed to manuscript revision.

## Data Availability

Example data and phasor analysis codes will be made available. Full datasets are available from the authors upon request.

## Declarations

A.J.B. is an inventor on patents PCT/US2019/062640; US 11,965,780; and US 11,828,851 describing wide-field, scanning, and resonant EO-FLIM.

---

**Extended Data Fig. 1.**
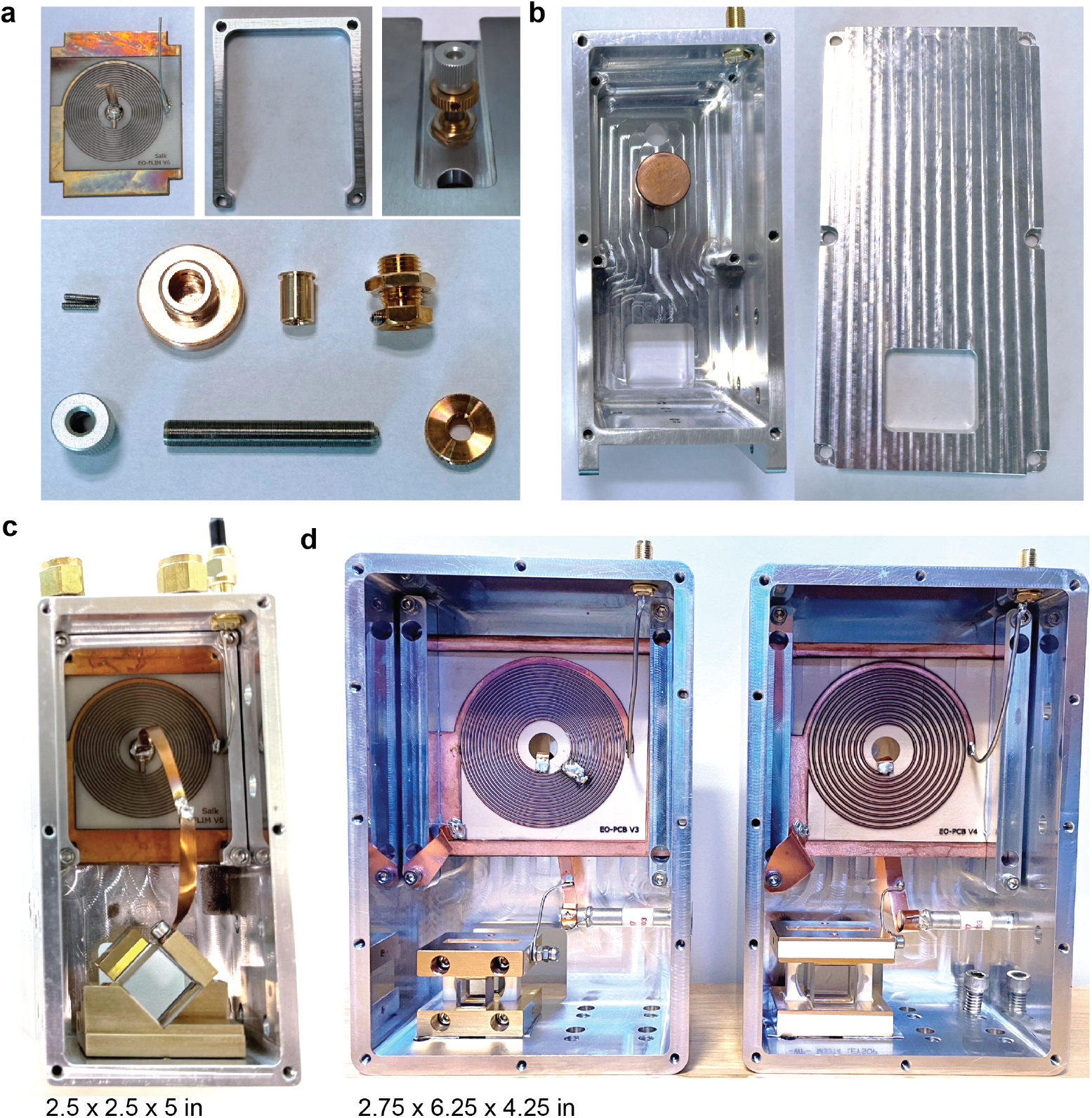
Compact planar resonant Pockels cells for EO-FLIM. Two designs were developed for 20 MHz RF drive, resulting in 40 MHz optical gating. The most compact design (panels **a**-**c**) uses an adjustable copper plunger behind the planar ceramic circuit as a trim capacitor for frequency tuning. The larger design (**d**) uses a commercial high voltage 10 pF trim capacitor (Sprague Goodman SGNMNC1103). The copper plunger has better thermal stability and lower RF loss. There are two Pockels cell mounting configurations compatible with each housing: one using a pair of lithium tantalate crystals rotated by 90 degrees with top electrodes connected by a copper foil bridge (panel c) and the other using aligned crystals and an achromatic half wave plate between them (panel d) (Leysop Inc.). Crystal surfaces have a single layer broadband AR coating. (**a**) Planar ceramic coil board (left), board clamp (middle), and thumbscrew adjuster (right) for the variable capacitor with exploded parts shown at bottom (Thorlabs 3/16”-100 thumbscrew and adapters — part numbers F19SS150, N100B2P, N100L5P, LN19100, and F19SSK1). (**b**) Aluminum housing and lid showing copper tuning plunger. A U-shaped channel through the back of the housing allows water cooling. RF input is through an SMA bulkhead connector. (**c**) Assembled compact resonant PC with 15 mm aperture crystals (90 degree rotated design). (**d**) Assembled larger design resonant PCs with 10 and 15 mm aperture crystals (half wave plate design). Optical gating performance is shown for both PC mounts in Supplementary Fig. 1d-g

**Extended Data Fig. 2.**
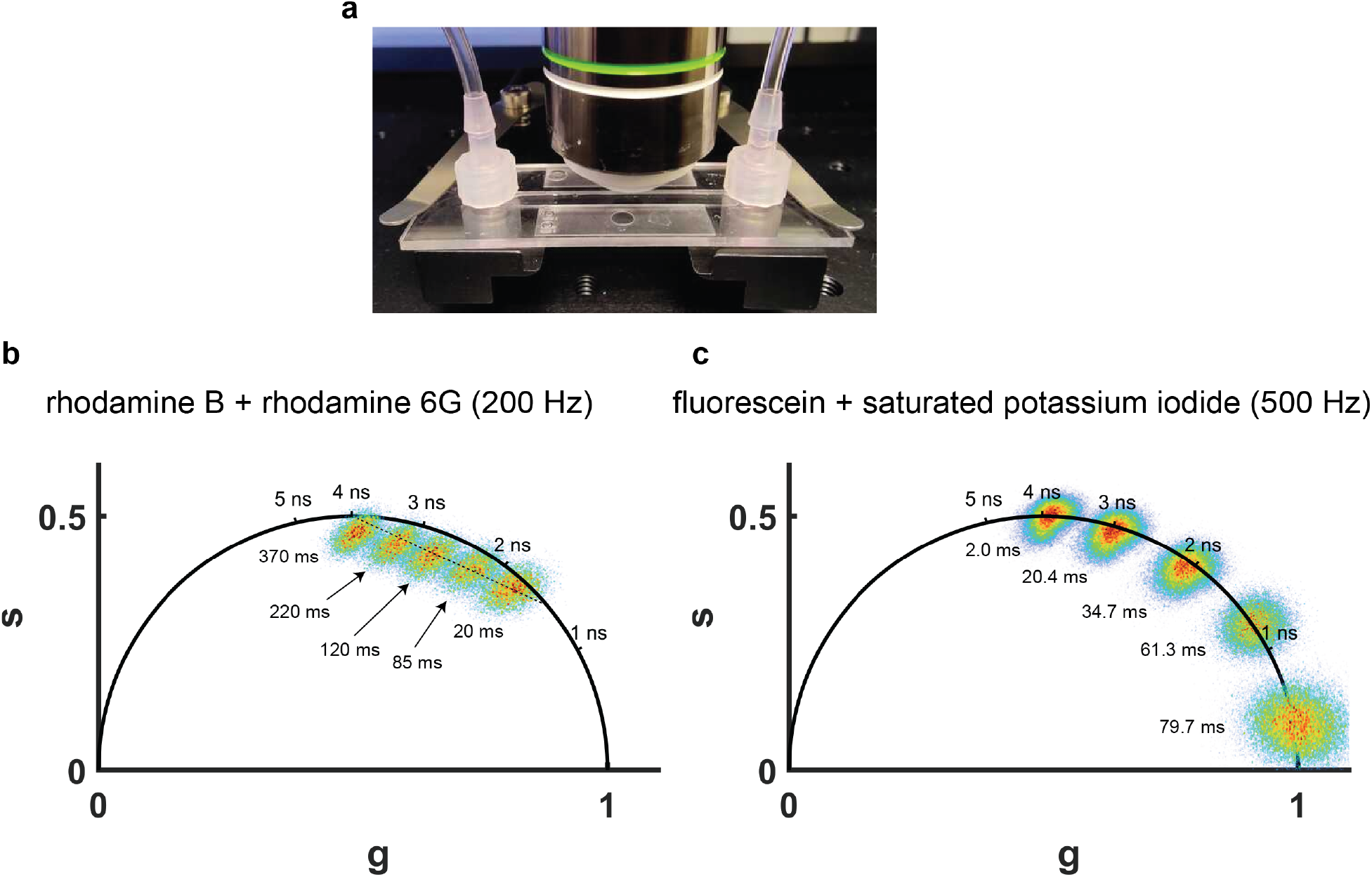
Fast phasor FLIM imaging of dye solution mixing and quenching. (**a**) Experimental setup for fast phasor FLIM imaging of dyes in a fluidic channel. (**b**) Phasor FLIM imaging at 200 Hz of rhodamine B mixed with rhodamine 6G, corresponding to Supplementary Video 1; representative phasor plots are shown at 20, 85, 120, 220, and 370 ms. (**c**) Phasor FLIM imaging at 500 Hz of fluorescein mixed with saturated potassium iodide quenched fluorescein, corresponding to Supplementary Video 2; representative phasor plots are shown at 2, 20.4, 34.7, 61.3, and 79.7 ms.

**Extended Data Fig. 3.**
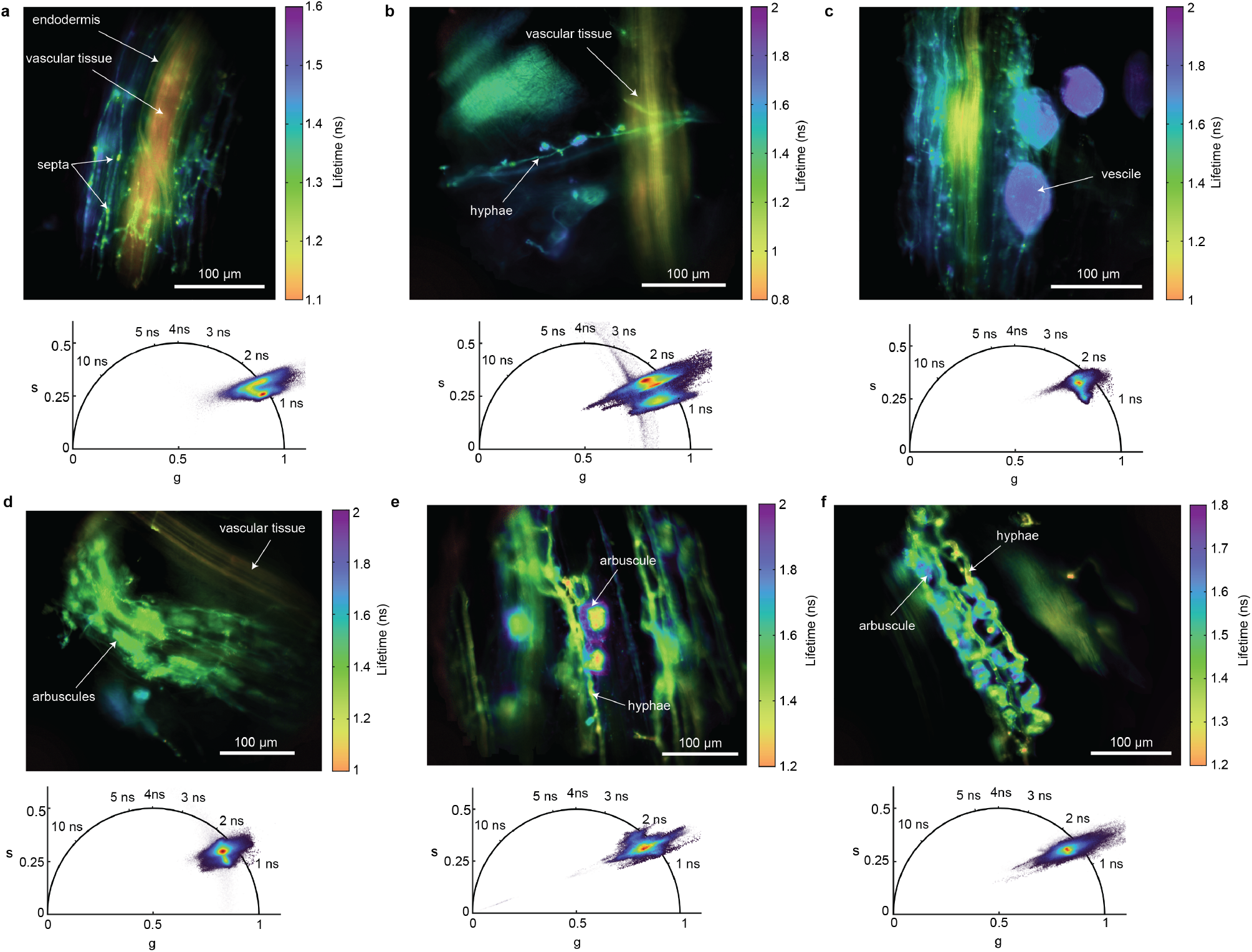
Phase lifetime images and corresponding phasor plots of cleared *Medicago truncatula* roots colonized by symbiotic *Rhizophagus irregularis* (**a**-**d**) *Paraglomus occultum* (**e**) and *Gigaspora rosea* (**f**) fungi stained using Alexa Fluor 488 WGA. Arrows and labels indicate fungal and root structures that are differentiated based on their lifetime, including fungal hyphae, vesicles, and arbuscules. Punctate septa are visible along hyphae segments with different lifetime characteristics. Root layers are also clearly differentiated in FLIM (a).

**Extended Data Fig. 4.**
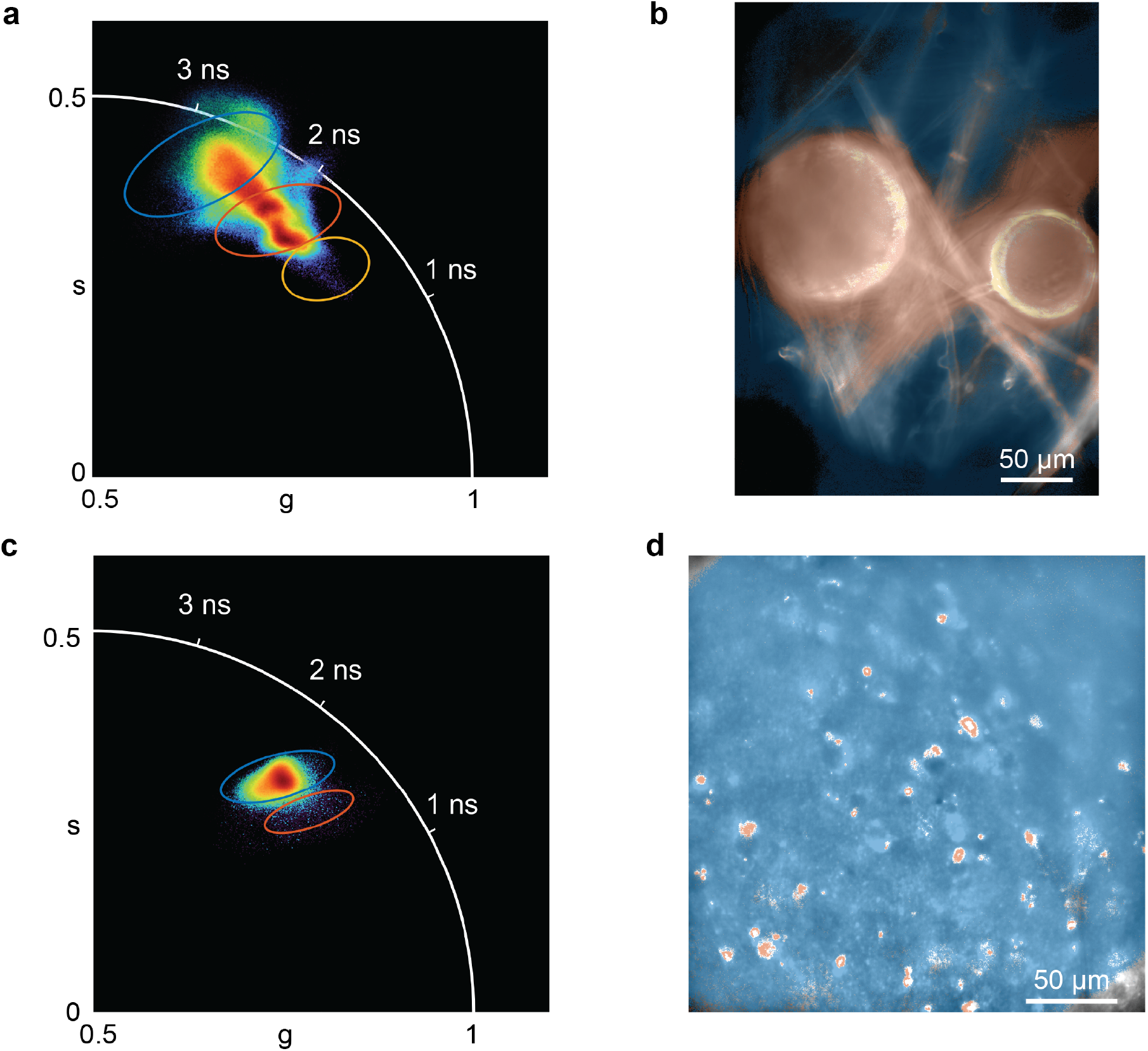
Image segmentation based on phasor coordinates. Image spatial structures at right are shown along with their associated color-coded phasor plot regions of interest at left. **(a**,**b)** *Rhizophagus irregularis* fungal spores from Fig. 1g, and **(c**,**d)** acute brain slice from Fig. 1h.

**Extended Data Fig. 5.**
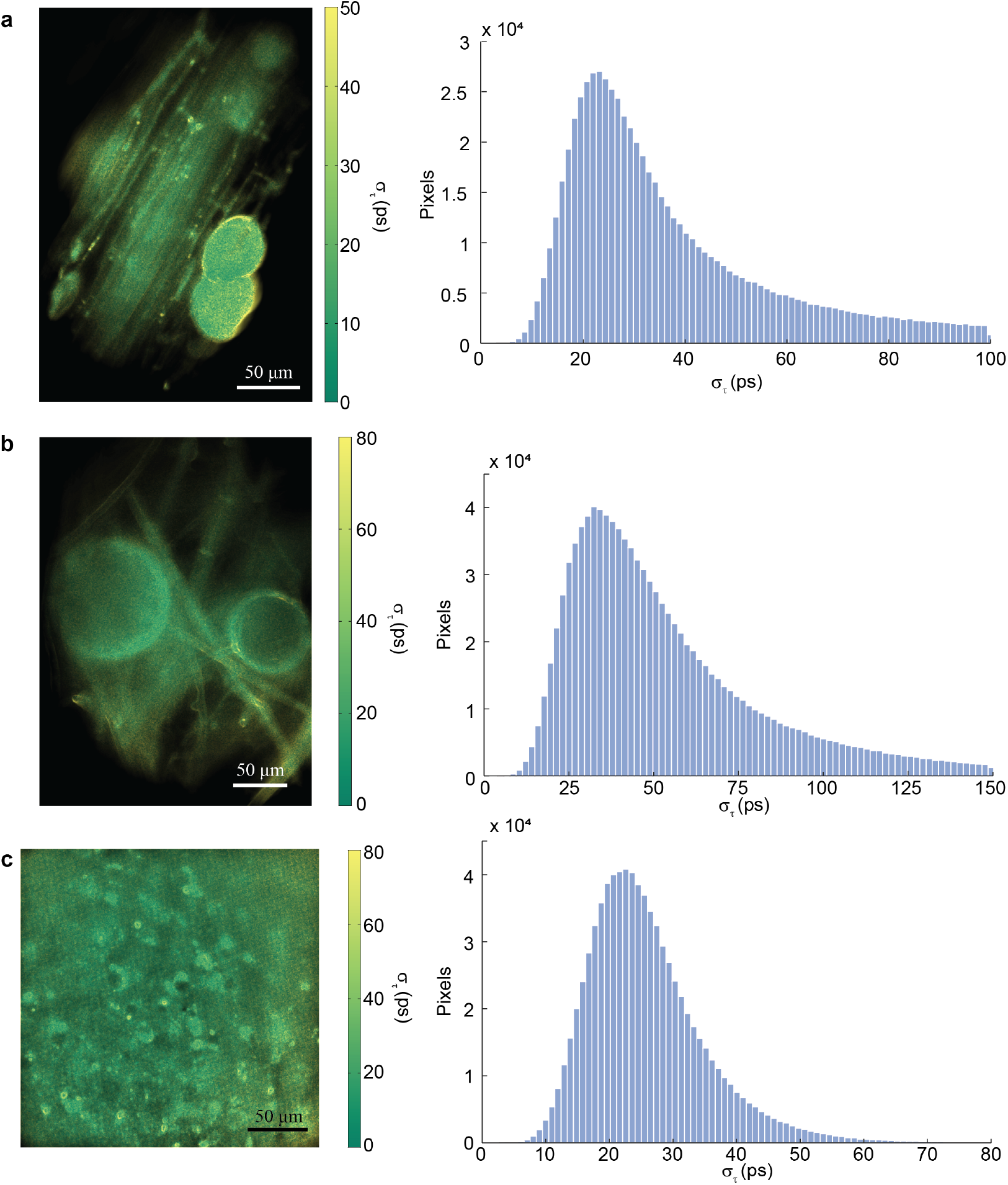
Local pixel precision for lifetime imaging. The standard deviation of the phase lifetime (*σ*_*τ*_) for each pixel was calculated from the 3 × 3 neighborhood centered on that pixel. (**a - c**) *σ*_*τ*_ image of *Medicago truncatula* root (Fig. 1e), *Rhizophagus irregularis* fungal spores (Fig. 1g), and acute brain slice (Fig. 1h) with their corresponding *σ*_*τ*_ histograms.

## Supplementary Information

**Supplementary Fig. 1.**
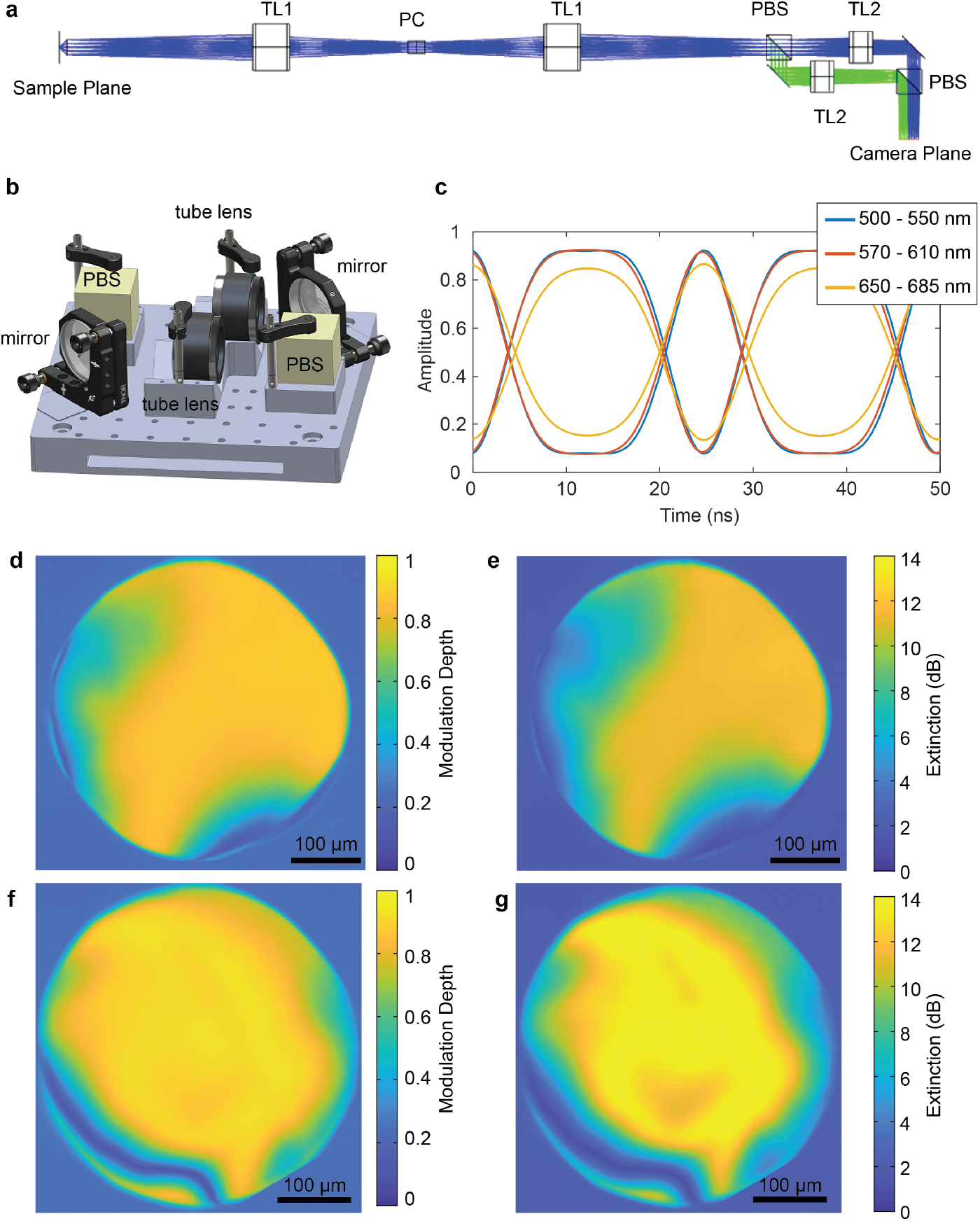
System design and gating characterization. (**a**) Optical schematic. Ray tracing was performed in Ansys Zemax OpticStudio. TL1: tube lens (Thorlabs, TTL200MP); TL2: tube lens (Thorlabs, TTL200); PC, Pockels cell; PBS, polarizing beam splitter (Bernhard Halle Nachfl., PTW 40). (**b**) Compact image splitter comprising two polarizing beam splitter cubes and two tube lenses. (**c**) Instrument response functions (IRFs) in three emission bands for a 10 mm PC crystal. The 500-550 nm IRF was measured using quenched fluorescein with an RF power of 25 W. The 575-610 nm IRF was measured using quenched fluorescein with an RF power of 30 W. The 650-685 nm IRF was measured using methylene blue with an RF power of 32 W. (**d**) Extinction is defined as 10 log_10_(max*/*min), where max and min are the extrema of the measured intensity trace in the gated channel for a 10 mm aperture Pockels cell (Extended Data Fig. 1d - waveplate design) (**e**) Modulation depth, calculated as (max − min)*/*(max + min) for the same Pockels cell. (**f**) Extinction for a 15 mm aperture Pockels cell (Extended Data Fig. 1c - 90 degree rotated design). (**g**) Modulation depth for the same Pockels cell.

**Supplementary Fig. 2.**
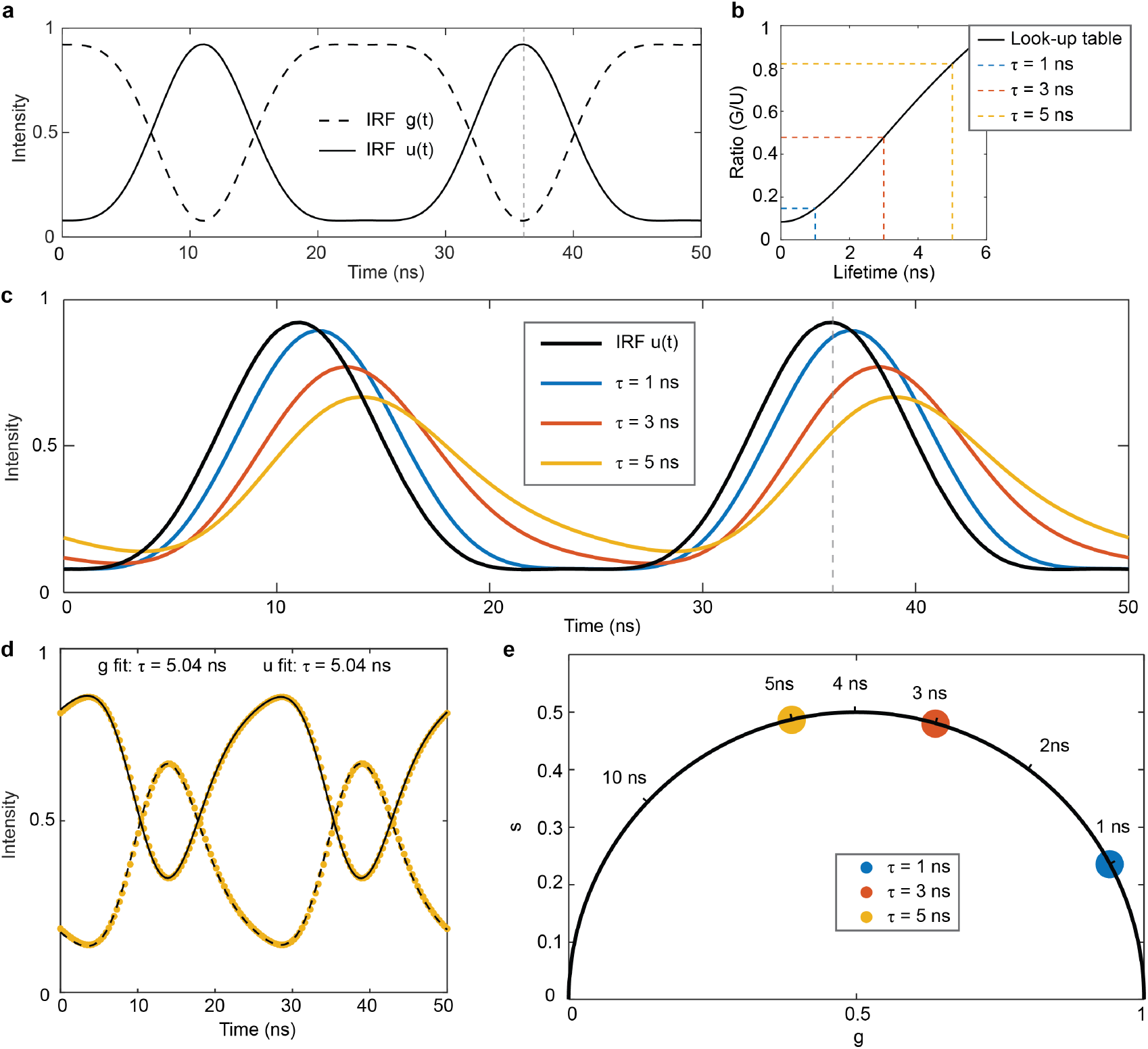
EO-FLIM principle and analysis methods. (**a**) Experimental instrument response functions for the gated channel, *g*(*t*), and the ungated channel, *u*(*t*). The dashed line indicates the optimal phase point used to generate the lookup table for single-phase acquisition. (**b**) Lookup table generated at the optimal phase, mapping the *G/U* ratio to the estimated lifetime. (**c**) Output *U* intensity traces in the ungated channel as a function of time for the IRF and for lifetimes of 1, 3, and 5 ns. The dashed line indicates the phase point used for the lookup table in (**b**). (**d**) Least squares fitting of the output intensity trace to extract the lifetime, shown for *τ* = 5 ns. (**e**) Phasor analysis maps lifetimes onto a 2D polar plot, where monoexponential decays lie on the universal semicircle and multiexponential decays appear as intensity-weighted geometric combinations of their components within the semicircle (see Supplementary Figs. 4-7).

**Supplementary Fig. 3.**
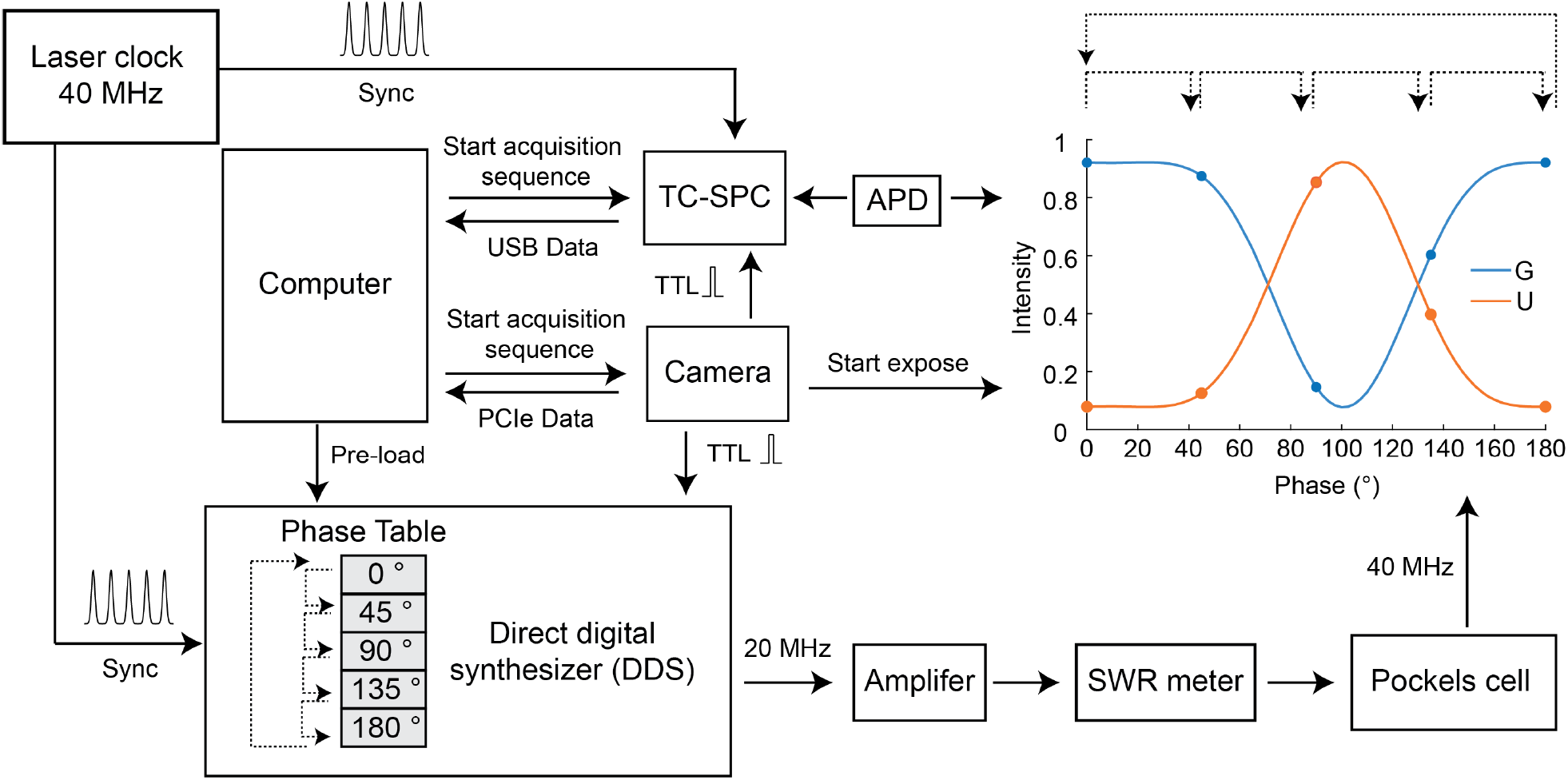
Electronic schematic. The pulsed laser provides the clock for synchronizing both the TC-SPC module and direct digital synthesizer (DDS). The DDS generates a 20 MHz sinusoidal drive with a programmable phase offset, which is subsequently amplified and applied to the resonant Pockels cell. For each FLIM acquisition, the computer initiates a synchronized acquisition sequence for the TC-SPC electronics and the camera. After each camera exposure, the camera outputs a trigger to both the DDS and the TC-SPC module. Upon receiving this trigger, the TC-SPC starts a new photon arrival histogram for the next frame exposure. In single-phase mode, the DDS ignores the frame trigger and maintains a constant programmed phase offset. In multi-phase mode, each frame trigger advances the DDS to the next phase point in a preloaded phase table. After the final entry, the DDS returns to the first phase point and is ready to begin the next phase cycle. Five phase points for rapid phasor acquisition are shown in the figure.

**Supplementary Fig. 4.**
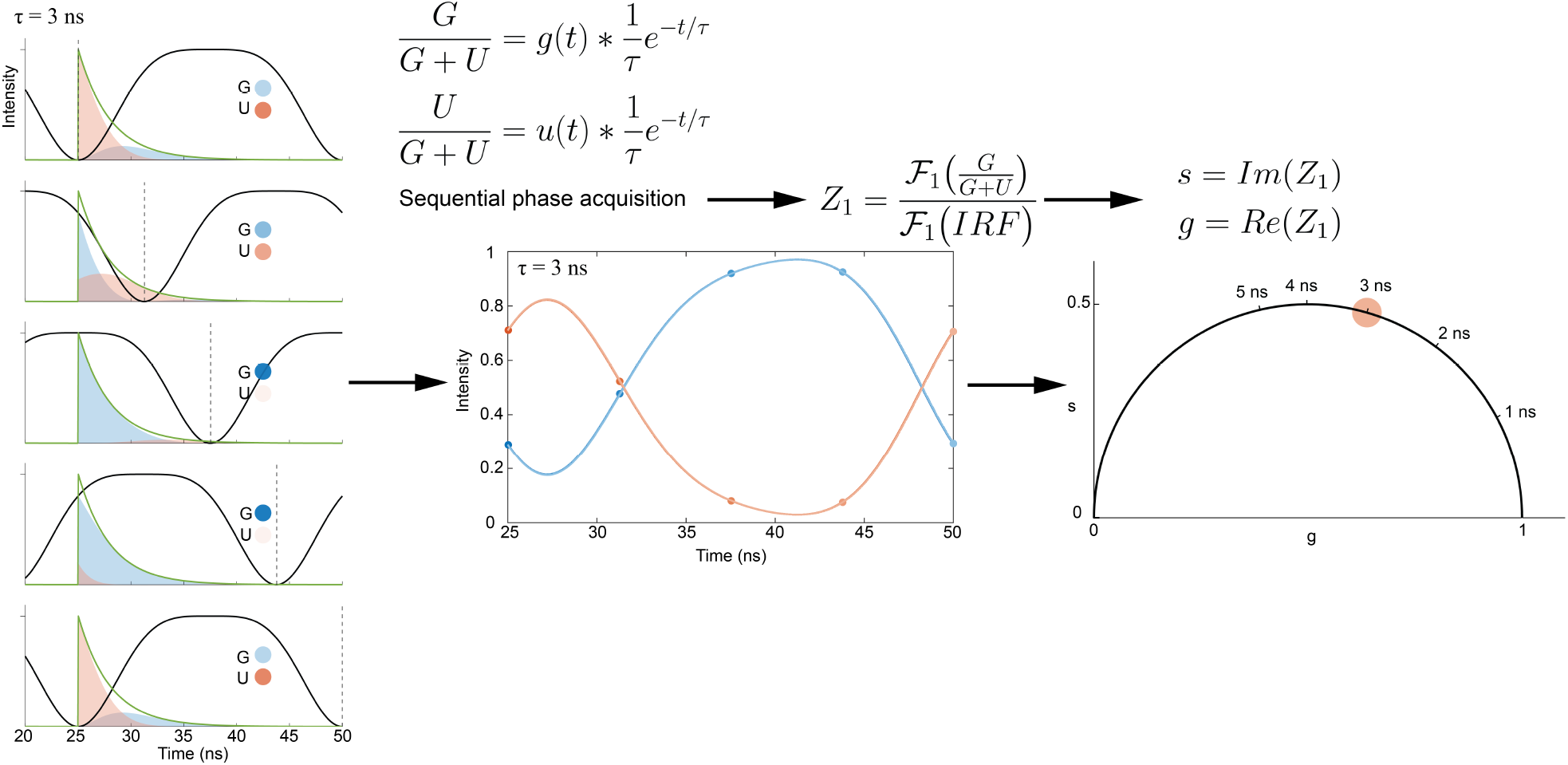
Rapid phasor FLIM acquisition using five phase points. A fluorescence lifetime decay (*τ* = 3 ns) is gated with the instrument response function (IRF) at five phase delays (left – 0°, 45°, 90°, 135°, and 180°). The captured points sample the convolution of the fluorescence decay with the gate (middle). The phasor coordinate (right) is calculated from the first harmonic Fourier coefficients after deconvolution with the IRF.

**Supplementary Fig. 5.**
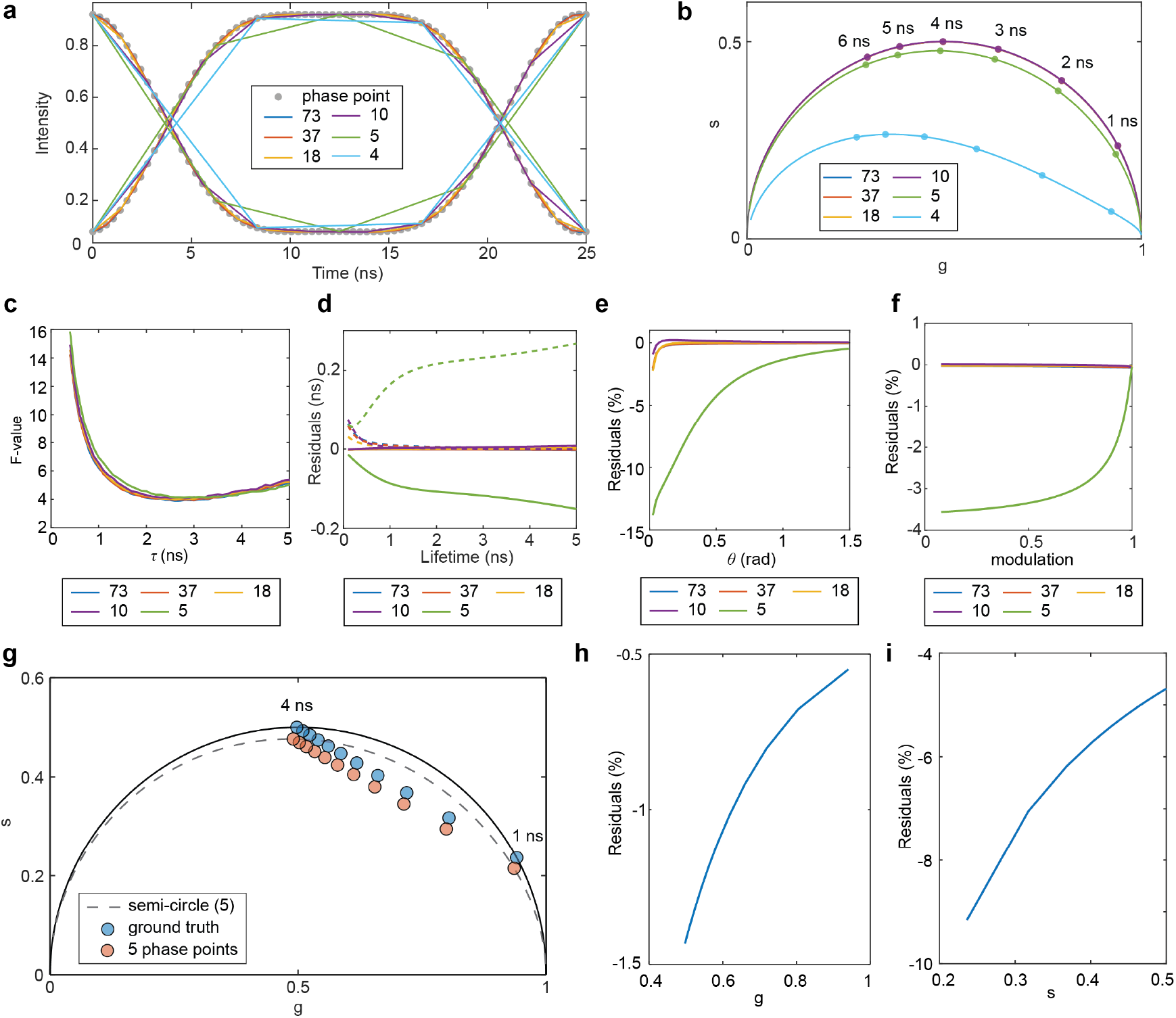
Effect of EO-FLIM phasor sampling. (**a**) Sub-sampled IRFs with different numbers of phase points (73, 37, 18, 10, 5, and 4) are plotted across one full gating cycle. (**b**) Phasor plots are constructed using different numbers of sampled phase points for single exponential lifetimes showing deviation from the universal semicircle. (**c**) F-values for EO-FLIM phasor analysis as a function of lifetime for different numbers of sampled phase points. (**d**) Corresponding lifetime residuals. Dashed lines indicate modulation lifetime, and solid lines indicate phase lifetime. (**e**) Phase angle residuals, and (**f**) modulation residuals. (**g**) Phasor plot of an example bi-exponential decay model (*τ*_1_ = 1 ns, *τ*_2_ = 4 ns) using five phase points at different mixture fractions *α*_1_ (*α*_1_ = 0, 0.1, 0.2, …, 1.0, with *α*_2_ = 1 *− α*_1_). (**h**) Residuals of *g* and (**i**) *s* for the bi-exponential decay in **g**.

**Supplementary Fig. 6.**
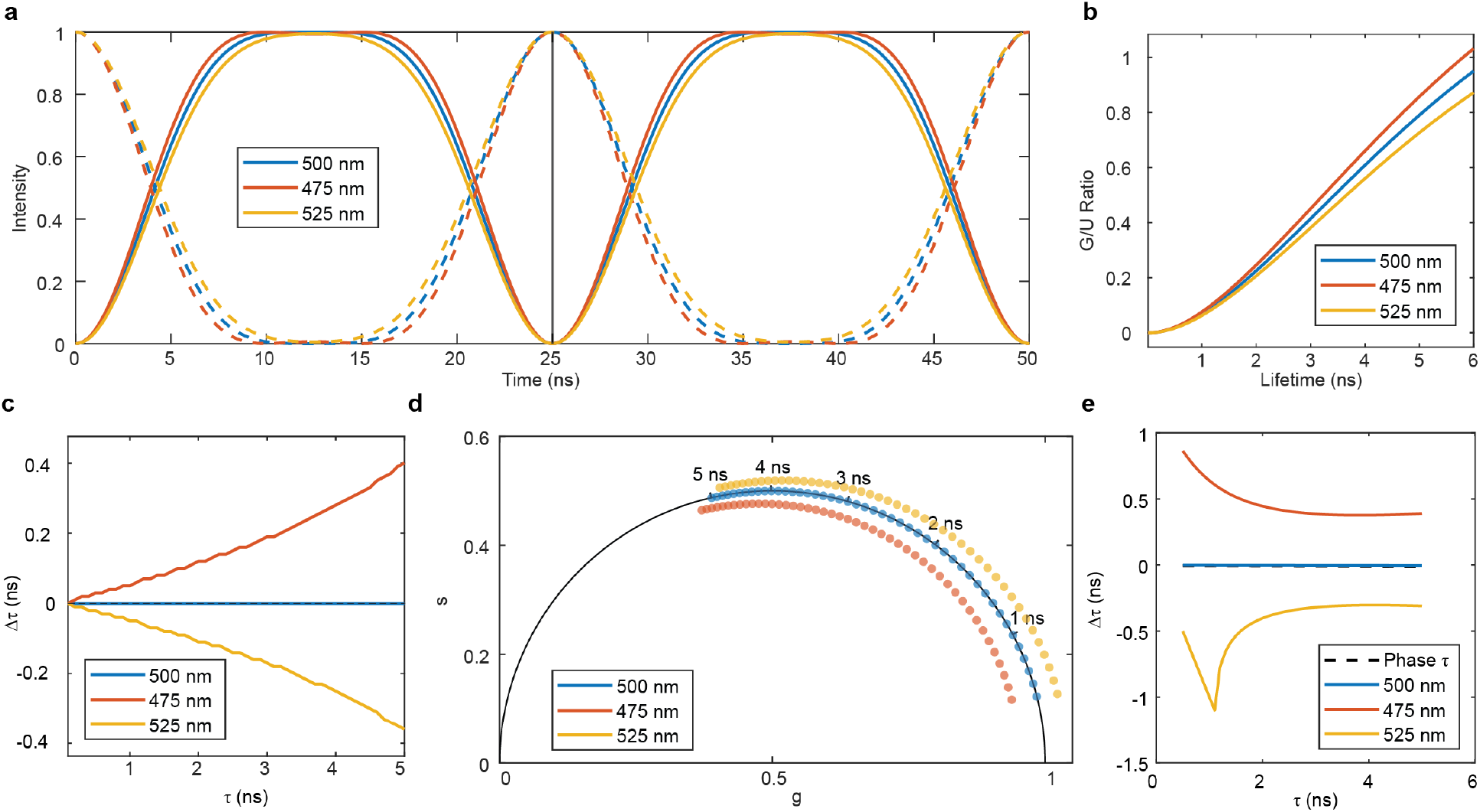
Effect of wavelength-dependent gating on EO-FLIM estimation. (**a**) EO-FLIM gating functions at the center wavelength (*λ* = 500 nm) and at the two band edges (475 and 525 nm). The gating function at the central wavelength is 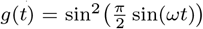, and the modified gating functions at the band edges are 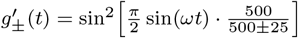. The black line indicates the hase point used to generate the lookup table. (**b**) Corresponding *G/U* ratio to lifetime lookup table generated at the indicated phase for single-phase acquisition. (**c**) Lifetime error versus ground-truth lifetime for 475, 500, and 525 nm using the *λ* = 500 nm lookup table. (**d**) Phasor plot for single-exponential decays at 475, 500, and 525 nm using the *λ* = 500 nm gating function. (**e**) Lifetime errors for 475, 500, and 525 nm. Phase lifetimes show no deviation and are constant across wavelengths (black dashed line) while modulation lifetimes show wavelength-dependent error (solid colored lines).

**Supplementary Fig. 7.**
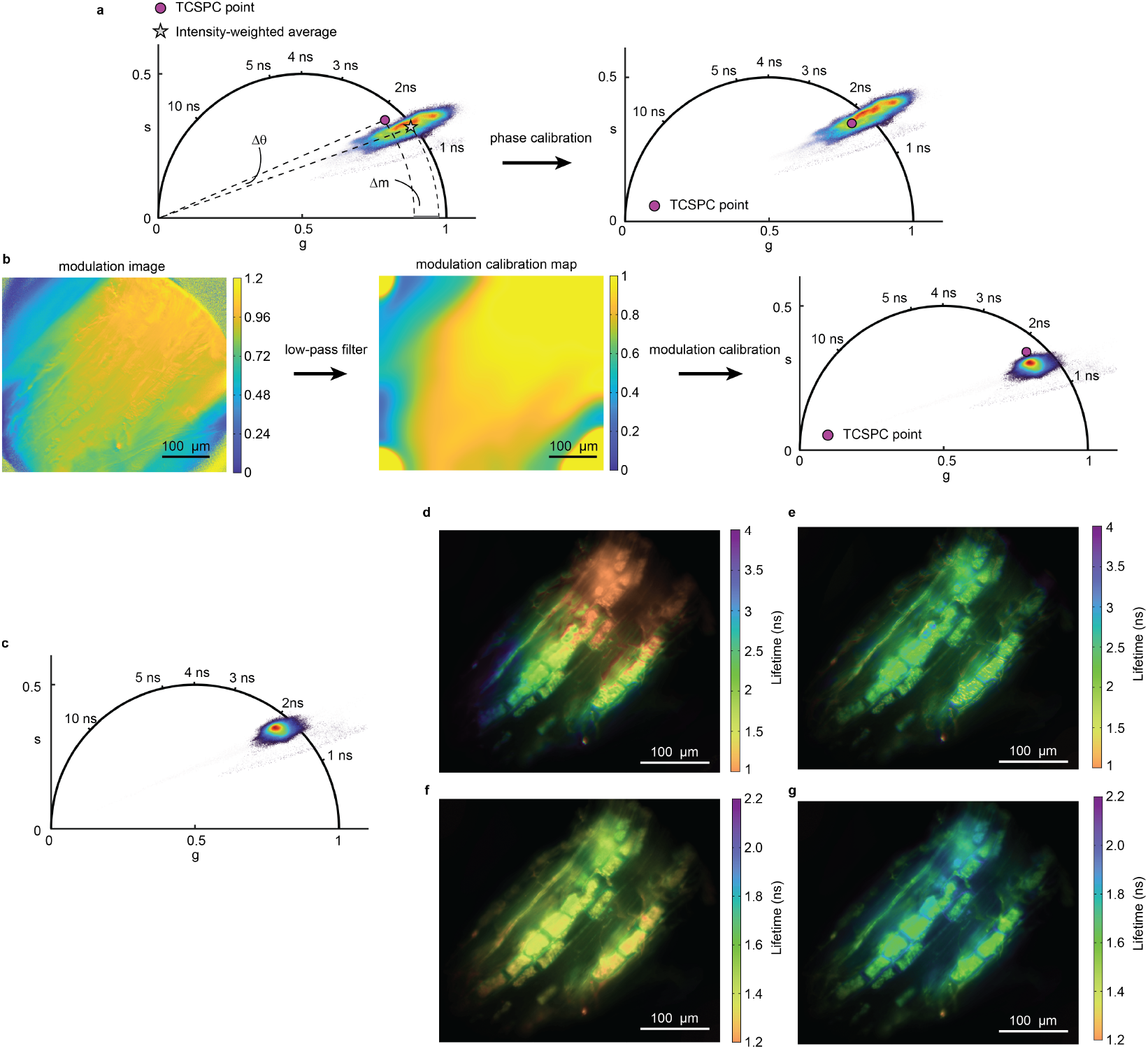
Phasor plot calibration with TC-SPC. The workflow for calibrating phase angle and modulation depth using spatially averaged TC-SPC data is illustrated with a representative *Medicago truncatula* root image. (**a**) Phase angle calibration is achieved by rotating all the coordinates by an angle Δ*θ*, where Δ*θ* is the phase angle difference between the TC-SPC reference point and the intensity-weighted average coordinates of all pixels. (**b**) For modulation depth calibration, the modulation image is low-pass filtered to obtain a low spatial frequency modulation image, which serves as the modulation calibration map. A modulation correction image is generated from the difference (Δ*m*) between the modulation calibration map and the TC-SPC coordinate and is applied to the modulation image. (**c**) Phasor plot after applying both phase and modulation calibrations. Note that typical phase corrections are small and not usually required (this is a larger example with Δ*θ* = 3.5° (280 ps correction), but typical Δ*θ* is < 1° (100 ps correction)). Modulation deviation results from crystal gating inhomogeneity as well as wavelength dependent gating shifts. (**d - g**) Lifetime images of the *Medicago truncatula* root calculated from (**d**) modulation depth, (**e**) phase angle, (**f**) calibrated modulation depth, and (**g**) calibrated phase angle.

**Supplementary Fig. 8.**
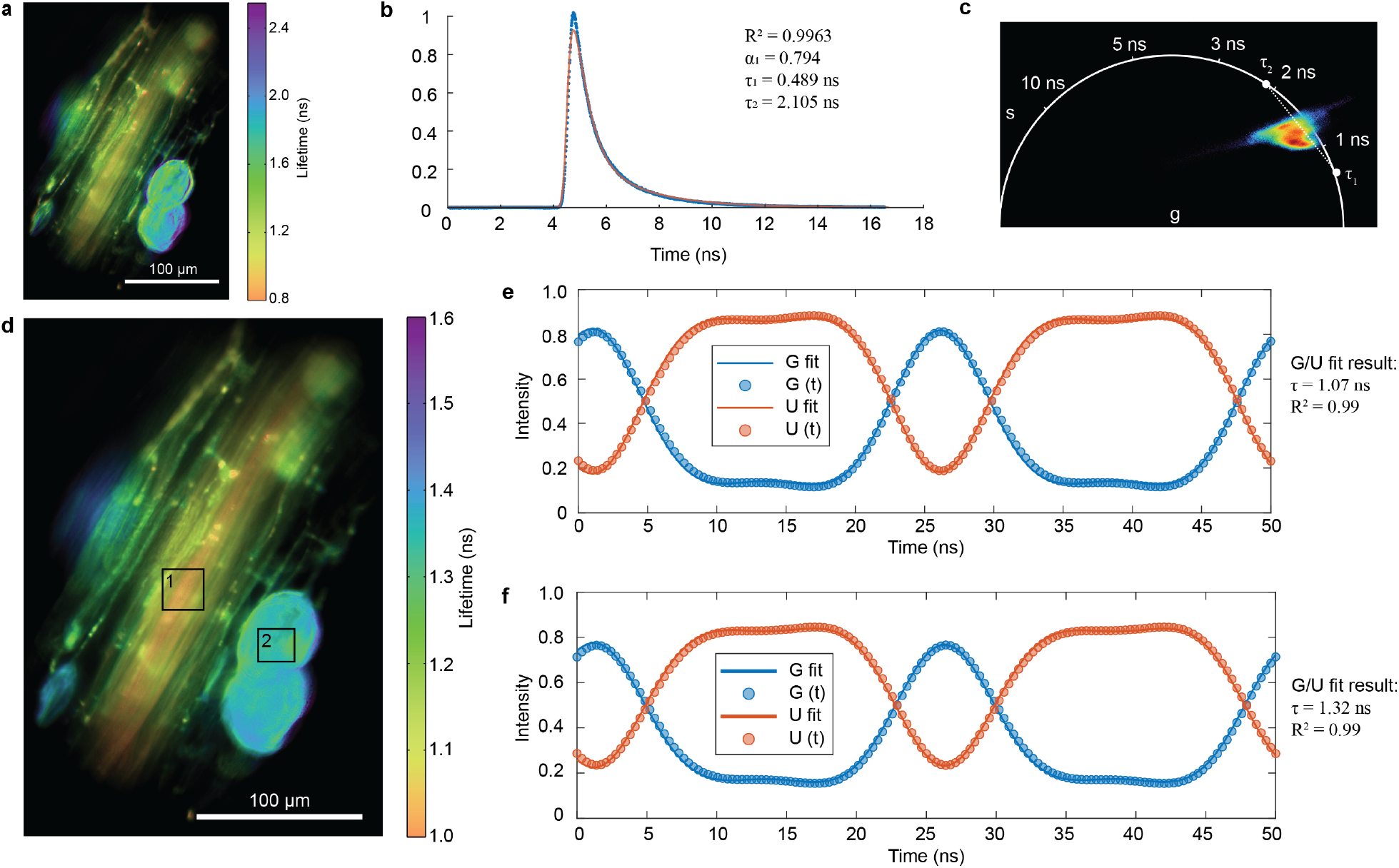
Lifetime estimation using alternative methods for the *Medicago truncatula* root image shown in Fig. 1e as phase lifetime. (**a**) Lifetime image obtained using single-phase analysis. (**b**) Spatially averaged TC-SPC histogram and bi-exponential fit. (**c**) Phasor plot with a reference line connecting *τ*_1_ and *τ*_2_, where *τ*_1_ and *τ*_2_ are derived from TC-SPC curve in (**b**). (**d**) Lifetime image obtained by single-exponential least squares fitting of the gated (*G*) intensity trace at each pixel. Single-exponential decay fitting for (**e**) ROI 1 and (**f**) ROI 2 indicated in (**d**).

## Supplementary Videos

**Supplementary Video 1** Phasor FLIM imaging at 200 Hz of rhodamine B mixing with rhodamine 6G, corresponding to Extended Data Fig. 2b. Two syringes of dye component solutions were exchanged through a fluidic channel.

**Supplementary Video 2** Phasor FLIM imaging at 500 Hz of fluorescein mixing with quenched fluorescein solution, corresponding to Extended Data Fig. 2c. Two syringes of dye and quenched dye solutions were exchanged through a fluidic channel.

**Supplementary Video 3** Phasor FLIM imaging of a crawling *Drosophila* larva at 40 Hz, corresponding to Fig. 2a,b. The phasor plot color and transparency were defined by the intensity.

**Supplementary Video 4** Phasor FLIM recording of sub-threshold voltage activity in a *Drosophila* neuron at 200 Hz, corresponding to Fig. 2c-g. (**a**) Δ*τ* image of the neuron. **(b)** Phasor plot of the image field shown in (**a**), where the red dot represents the average phasor coordinates of the circled ROI. **(c)** Mean *τ* over time calculated within the ROI.

**Supplementary Video 5** Phasor FLIM recording of action potentials in a *Drosophila* neuron at 500 Hz, corresponding to Fig. 2h. **(a)** Phase lifetime image of the neuron (scale bar = 8 *µm*) and corresponding phasor plot, where the red dot indicates the average phasor coordinates of the ROI marked in (**a**). **(b)** Mean intensity over time calculated within the ROI. **(c)** Mean phase lifetime over time calculated within the ROI.

### EO-FLIM Derivations

#### Single-phase EO-FLIM

The EO-FLIM resonant gating function is

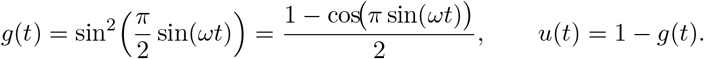

Using the Jacobi–Anger expansion with optical gating frequency Ω equal to twice the RF drive frequency (Ω = 2*ω*), we can write the real Fourier series

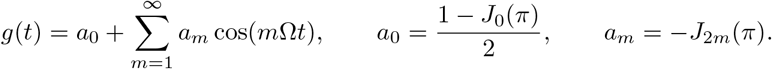

For a single-exponential decay 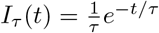, the normalized gated signal is

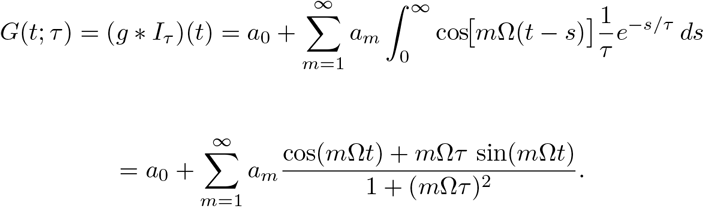

The ratio-to-lifetime lookup table is

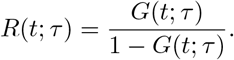

#### Phasor EO-FLIM

For deriving phasor EO-FLIM we use the equivalent complex Fourier series coefficients

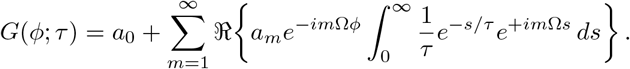

The coefficient of the *m*-th harmonic after convolution is

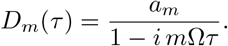

Normalizing by the zero-lifetime instrument response function *D*_*m*_(0) = *a*_*m*_ gives

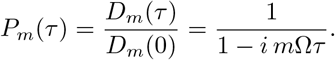

The g and s phasor coordinates correspond to the real and imaginary components

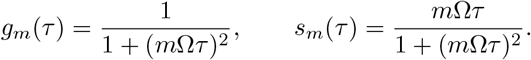

For the first harmonic,

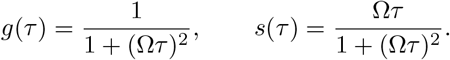

The modulation and phase are given by

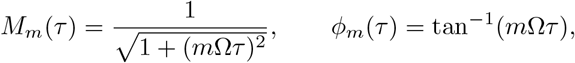

and the corresponding lifetimes are the standard expressions for phasor FLIM

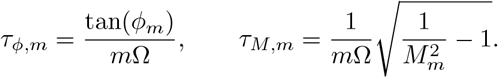

#### Wavelength-dependent gating

If the half-wave voltage is set for wavelength *λ*_0_, the wavelength-dependent gate becomes

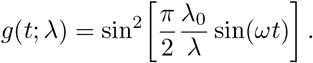

Resulting in wavelength-dependent Fourier coefficients

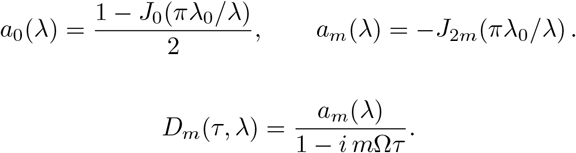

Since *a*_*m*_(*λ*) is real, wavelength shifts affect the modulation but not the phase coordinate (Supplementary Fig. 7d). The lookup table for single-phase estimation is modified through *G*(*t*; *τ, λ*) (Supplementary Fig. 7b). Note that this effect may be exploited for multi-dimensional measurements of spectrum and lifetime, for example by adding a quarter waveplate in front of the Pockels cell to maximize wavelength sensitivity [9].

## References

[1] Digman, M. A., Caiolfa, V. R., Zamai, M. & Gratton, E. The Phasor Approach to Fluorescence Lifetime Imaging Analysis. Biophysical Journal 94, L14–L16 (2008).

[2] Esposito, A., Oggier, T., Gerritsen, H. C., Lustenberger, F. & Wouters, F. S. All-solid-state lock-in imaging for wide-field fluorescence lifetime sensing. Optics Express 13, 9812–9821 (2005).

[3] Chen, H., Holst, G. & Gratton, E. Modulated CMOS camera for fluorescence lifetime microscopy. Microscopy Research and Technique 78, 1075–1081 (2015).

[4] Raspe, M. et al. siFLIM: single-image frequency-domain FLIM provides fast and photon-efficient lifetime data. Nature Methods 13, 501–504 (2016).

[5] Sorrells, J. E. et al. Real-time pixelwise phasor analysis for video-rate two-photon fluorescence lifetime imaging microscopy. Biomedical Optics Express 12, 4003–4019 (2021).

[6] Zhang, Y. et al. Instant FLIM enables 4D in vivo lifetime imaging of intact and injured zebrafish and mouse brains. Optica 8, 885–897 (2021).

[7] Ulku, A. et al. Wide-field time-gated SPAD imager for phasor-based FLIM applications. Methods and Applications in Fluorescence 8, 024002 (2020).

[8] Dunsing-Eichenauer, V. et al. Fast volumetric fluorescence lifetime imaging of multicellular systems using single-objective light-sheet microscopy. Communications Biology 8, 1785 (2025).

[9] Bowman, A. J., Klopfer, B. B., Juffmann, T. & Kasevich, M. A. Electrooptic imaging enables efficient wide-field fluorescence lifetime microscopy. Nature Communications 10, 4561 (2019).

[10] Bowman, A. J. & Kasevich, M. A. Resonant Electro-Optic Imaging for Microscopy at Nanosecond Resolution. ACS Nano 15, 16043–16054 (2021).

[11] Bowman, A. J., Huang, C., Schnitzer, M. J. & Kasevich, M. A. Wide-field flu-orescence lifetime imaging of neuron spiking and subthreshold activity in vivo. Science 380, 1270–1275 (2023).

[12] Marchand, R. et al. Super-resolution Live-cell Fluorescence Lifetime Imaging. Preprint at http://arxiv.org/abs/2502.16672 (2025).

[13] Knight, V. R. et al. Fast wide-field light sheet electro-optic FLIM. Optics Express 34, 15065–15073 (2026).

[14] Beauvais, A. & Latgé, J.-P. Special Issue: Fungal Cell Wall. Journal of Fungi 4, 91 (2018).

[15] Lamon, G. et al. Solid-state NMR molecular snapshots of Aspergillus fumigatus cell wall architecture during a conidial morphotype transition. Proceedings of the National Academy of Sciences 120, e2212003120 (2023).

[16] Kannan, M. et al. Dual-polarity voltage imaging of the concurrent dynamics of multiple neuron types. Science 378, eabm8797 (2022).

[17] Orosz, J. et al. The pseudokinase CORYNE modulates Medicago truncatula inflorescence meristem branching and plays a conserved role in the regulation of arbuscular mycorrhizal symbiosis. Journal of Experimental Botany 76, 7086–7104 (2025).

[18] Aso, Y. et al. The neuronal architecture of the mushroom body provides a logic for associative learning. eLife 3, e04577 (2014).

